# DeepFace: A High-Precision and Scalable Deep Learning Pipeline for Predicting Large-Scale Brain Activity from Facial Dynamics in Mice

**DOI:** 10.1101/2025.06.10.658952

**Authors:** Kemal Ozdemirli, Tenesha Connor, Kaleb Kim, Macit Emre Lacin, Miguel Maldonado, Thiago Peixoto Leal, Caglar Oksuz, Ignacio Mata, Murat Yildirim

**Affiliations:** Department of Neuroscience, Cleveland Clinic Lerner Research Institute, Cleveland, OH, USA; Department of Mechanical and Aerospace Engineering, Case Western Reserve University, Cleveland, OH, USA; Department of Biomedical and Chemical Engineering, Cleveland State University, Cleveland, OH, USA; Department of Computer and Data Sciences, Case Western Reserve University, Cleveland, OH, USA; Genomic Medicine Institute, Cleveland Clinic Lerner Research Institute, Cleveland, OH, USA; Comprehensive Cancer Center, Case Western Reserve University, Cleveland, OH, USA

## Abstract

We present DeepFace, a next-generation facial analysis pipeline that enhances orofacial tracking and cortical activity prediction in mice. Rather than replacing existing tools, DeepFace builds upon DeepLabCut and Facemap to address scalability bottlenecks and improve behavioral quantification. It offers high precision, keypoint customization, and robust performance across GCaMP6s, GCaMP6f, and jGCaMP8m lines. With scalable batch processing and high-performance computing compatibility, DeepFace enables high-throughput brain-behavior analysis in large-scale preclinical neuroscience.

## Main

Facial dynamics in both mice and humans are tightly coupled with brain activity. Spontaneous facial movements are embedded in widespread cortical signals, where neural activity often reflects internal behavioral states more strongly than responses to cognitive tasks^1,2^. High-dimensional neural data across multiple brain regions reflect these facial movements, redefining trial-to-trial variability as meaningful behavior-driven fluctuations^1^. Behavioral state also reorganizes functional connectivity in the cortex, further supporting the link between facial activity and brain-wide dynamics^3^. In humans, facial expressions are encoded in specific cortical regions, such as the posterior superior temporal sulcus^4^, while stimulation of sensory regions like the auditory cortex can evoke facial movements^5^, underscoring the integrative nature of sensorimotor networks. Facial analysis is emerging as a non-invasive tool for both disease detection and behavioral classification. In humans, facial recognition algorithms have been shown to detect genetic syndromes^6^ as well as identify Parkinson’s disease^7^, offering new avenues for early detection. In preclinical models, deep learning has been used to distinguish emotional states^8^ and identify facial motor deficits like facial paralysis in awake mice^9^.

Existing tools such as Facemap^10^ and DeepLabCut^11^ have enabled major advances in facial tracking and behavior analysis, yet large-scale applications can be limited by requirements for manual refinement or task-specific customization. To address these scalability and generalization challenges, we developed DeepFace, a high-precision facial tracking pipeline that integrates SLEAP^12^ for accurate, flexible pose estimation. DeepFace supports local and high-performance computing (HPC) deployment for ultrafast processing of large video datasets. Furthermore, we extend DeepFace with a deep learning model that robustly predicts cortical activity from facial features, enabling a new class of models linking external behavior to internal brain dynamics. This innovation provides a powerful and efficient platform for linking external facial behavior to internal brain dynamics, paving the way for robust, reproducible, and scalable disease classification models in both preclinical and clinical neuroscience.

First, we illustrate the DeepFace and DeepLabCut (DLC) workflow, which begins by extracting 330 representative frames through our **custom-built mediaGUI** from each of 90 behavioral videos spanning GCaMP6s, GCaMP6f, and jGCaMP8m lines (**Fig.1A, see Methods**). These frames were manually labeled for 12 facial keypoints, including eyelids, whiskers, nose, and mouth features. The labeled images were used to train a neural network—UNet for DeepFace and ResNet50 for DLC—which was then applied to the full-length videos for automated orofacial feature tracking through our custom-built SLEAP-GUI (**Supp. Fig.1A, see Methods**). Then, we present the Facemap workflow, which includes a base model with optional refinement through manual annotation and retraining (**Fig. 1B**). Across 90 video datasets (5 mice and 6 experiments for each mouse line), DeepFace consistently achieved higher precision in detecting orofacial features including the eyelid, nose, mouth, and whiskers. While tools like Facemap are highly useful and have enabled widespread applications in behavioral analysis, their refinement process can require manual effort for optimal performance in diverse contexts (**Supp. Fig. 1B**). In contrast, DeepFace generalized well across genotypes and experimental conditions, offering robust results with minimal user intervention. Facemap currently supports a fixed set of keypoints, optimized for common facial landmarks. DeepFace adds flexibility by enabling full customization of keypoint definitions, allowing users to tailor facial tracking to their specific experimental needs (**Supp. Fig. 2**). In precision analyses (**Fig. 1C**), DeepFace outperformed Facemap’s refined and base models as well as DeepLabCut in all individual mice as well as individual genotypic groups ((**GCaMP6s**: p < 0.05 vs Refined, p<0.0001 vs DLC and Basemodel, **GCaMP6f**: p < 0.001 vs. Refined, p < 0.0001 vs. DLC and Basemodel, **jGCaMP8m**: p < 0.001 vs. Refined and Basemodel, p < 0.0001 vs. DLC, paired t-test) (**Supp. Fig. 3, see Videos1-3**). Importantly, DeepFace’s performance remained stable across repeated experiments, indicating strong generalization and minimal overfitting. Beyond accuracy, DeepFace demonstrated substantial computational advantages, processing large datasets approximately an order of magnitude faster than DeepLabCut and Facemap’s refined models when deployed on high-performance computing (HPC) clusters (**Fig. 1D, see Methods**), highlighting its scalability and efficiency for large-scale analyses. Notably, analyzing 90 videos took approximately 13 hours with Facemap’s manual refinement, while DeepFace completed the same task in just 2.5 hours on an HPC system. This efficiency scales dramatically: processing 1,000 videos with DeepFace can be completed within a single day, fully automated—whereas Facemap would require nearly a month of manual intervention, making DeepFace a truly scalable solution for high-throughput behavioral analysis. These results underscore DeepFace’s potential as a scalable and efficient solution for high-throughput behavioral analysis. By combining accuracy, speed, and flexibility in a single pipeline, DeepFace supports reproducible and automated facial tracking across a wide range of experimental paradigms. Rather than replacing existing tools, DeepFace builds upon their foundations to address scalability bottlenecks and broaden the accessibility of detailed behavioral quantification for large-scale neuroscience studies.

**Figure 1.**
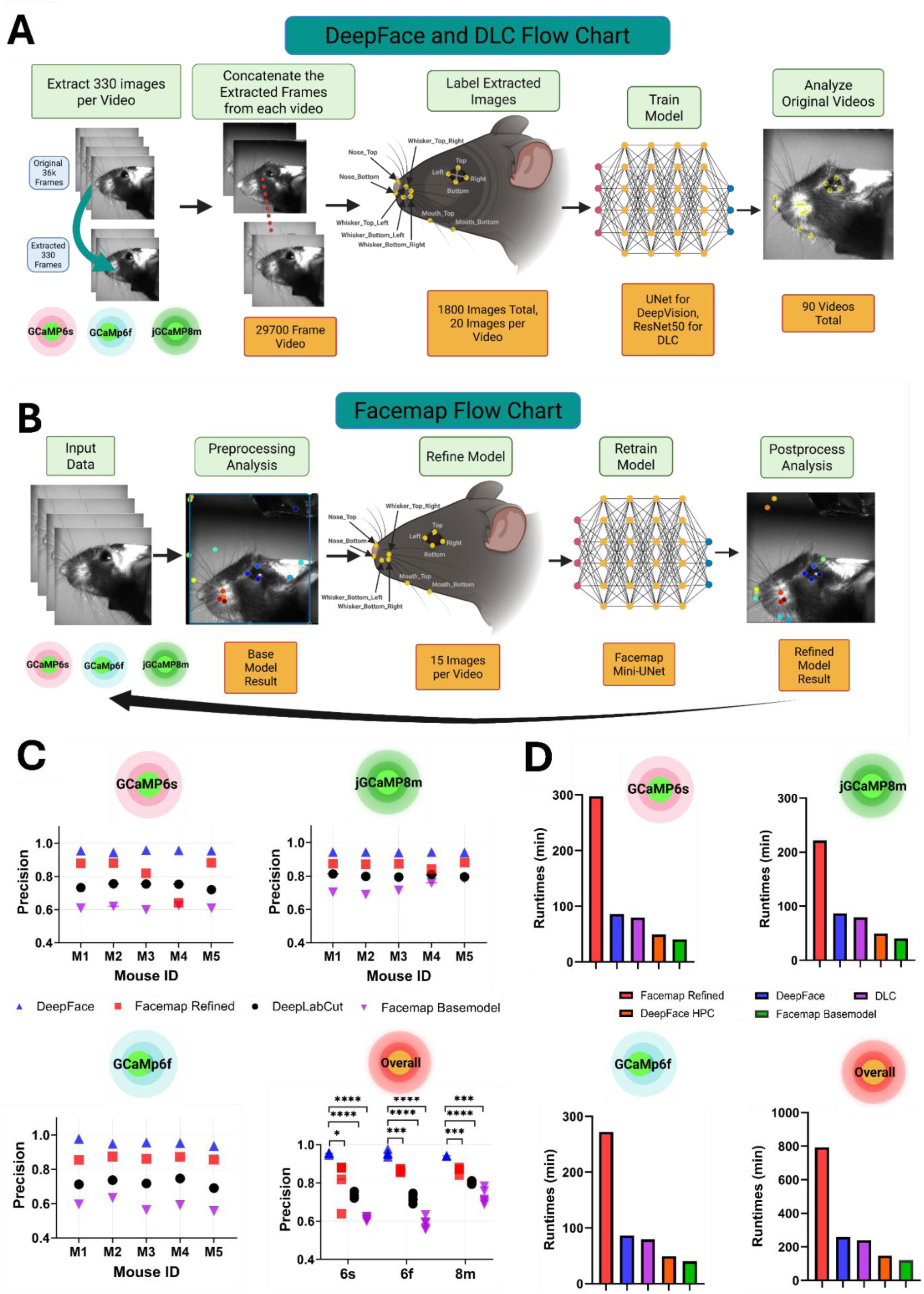
High-Precision and Ultrafast Orofacial Tracking Using DeepFace Across Multiple Transgenic Mouse Lines. (A) DeepFace and DeepLabCut Analysis Pipeline. To achieve robust and generalizable tracking of orofacial features across different calcium indicator mouse lines (GCaMP6s, GCaMP6f, and jGCaMP8m), we developed a standardized and scalable workflow. From each video (n = 90), 330 frames were uniformly extracted and concatenated into a single training video using a custom-built GUI. A total of 1800 frames were labeled with 12 keypoints (eye, nose, mouth, whiskers) and used to train models using both SLEAP (U-Net backbone) and DeepLabCut (ResNet-50). The trained model was then applied back to individual original videos for inference and benchmarking. (B) Facemap Refinement Workflow. In parallel, we evaluated the performance of the existing Facemap pipeline. Each video was first processed using the base Facemap model. Then, a minimal set of 15 frames per video was manually labeled with 11 keypoints, and the model was retrained using the Facemap Mini-UNet. This refinement loop was repeated for each video within a transgenic line, enabling adaptive fine-tuning across datasets. (C) Model Precision Across Transgenic Lines. Precision performance (mean ± SEM) was benchmarked for each of the 5 mice (M1–M5) across six experimental days and three transgenic lines. DeepFace consistently outperformed Facemap Base Model, Refined Facemap, and DeepLabCut in precision across GCaMP6s, GCaMP6f, and jGCaMP8m lines. The final panel aggregates performance across all lines, highlighting DeepFace’s superior and generalizable accuracy (GCaMP6s: p < 0.05 vs Refined, p<0.0001 vs DLC and Basemodel, GCaMP6f: p < 0.001 vs. Refined, p < 0.0001 vs. DLC and Basemodel, jGCaMP8m: p < 0.001 vs. Refined and Basemodel, p < 0.0001 vs. DLC, paired t-test). (D) Runtime Efficiency of Analysis Pipelines. We quantified the total processing time (in minutes) for each model across all videos and transgenic lines. DeepFace achieved high precision with minimal runtime overhead, significantly outperforming DeepLabCut and refined Facemap pipelines. DeepFace on a high-performance computing cluster (HPC) further reduced runtime, demonstrating scalability for large datasets.

To investigate whether orofacial dynamics can reliably predict brain-wide activity, we implemented a multimodal pipeline that synchronizes high-speed facial videography with widefield calcium imaging in head-fixed mice (**Fig. 2A**). During each 300-second session, facial videos were recorded at 120 Hz using an IR-illuminated camera, while brain activity was simultaneously captured using a custom-made 1-photon imaging system (**see Methods and Videos4-6**). For GCaMP6s and GCaMP6f mice^13^, blue and violet light alternated at 40 Hz to enable hemodynamic correction, producing corrected calcium signals at 20 Hz. For jGCaMP8m mice^14^, continuous blue light acquisition at 100 Hz was used to match the sensor’s fast kinetics. Brain recordings were spatially registered to the **Allen Brain Atlas**, enabling consistent extraction of activity from **11 cortical regions per hemisphere**, including motor, somatosensory, retrosplenial, and visual areas (**see Methods**). Facial videos were independently processed using three pose estimation pipelines: our DeepFace model, Facemap Basemodel and Facemap Refined model. Each method extracted the x– and y-coordinates of 12 keypoints representing eyelid, nose, mouth, and whisker positions. These coordinates were used to train neural networks for brain activity prediction, allowing us to directly compare how each facial tracking method performs in decoding large-scale cortical dynamics (**Supp. Fig. 4**).

**Figure 2.**
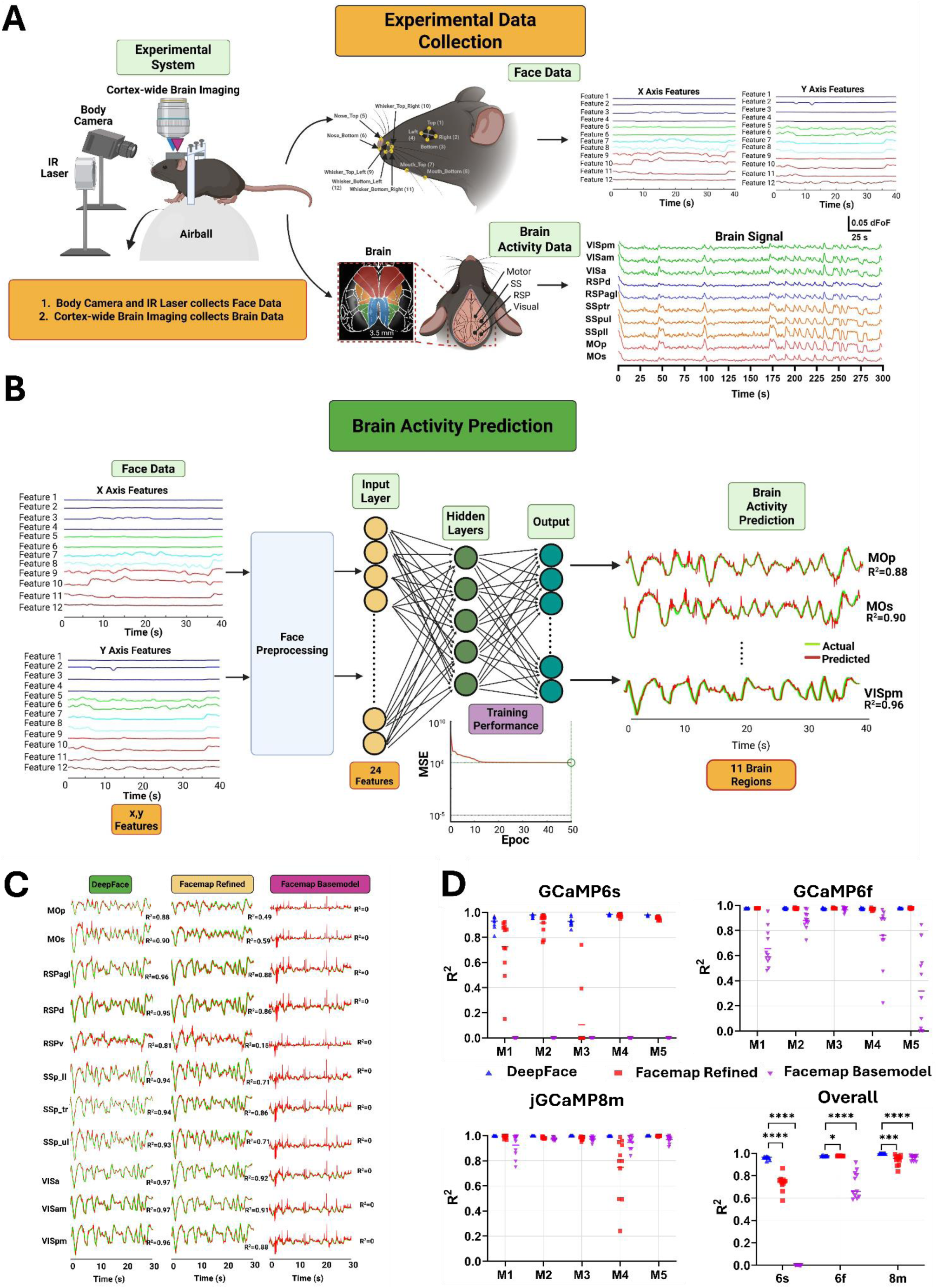
Decoding Brain Activity from Orofacial Dynamics Using DeepFace in Head-Fixed Mice. (A) Experimental Setup and Data Collection. We developed a high-resolution behavioral and neural data acquisition system combining 120 Hz facial videography and widefield one-photon calcium imaging to record from 11 cortical brain regions in both hemispheres. Three transgenic mouse lines (GCaMP6s, GCaMP6f, jGCaMP8m) underwent 6-day experiments, each lasting 5 minutes per session. Brain imaging was performed at 20 Hz (GCaMP6s, GCaMP6f) or 100 Hz (jGCaMP8m). Following each session, facial videos were processed through three pipelines—DeepFace, Facemap Refined, and Facemap Basemodel—extracting 12 facial features (x/y positions of eyelids, nose, mouth, and whiskers). Brain signals were segregated into 11 anatomically defined regions for downstream analysis. **(B) Deep Learning Model for Brain Activity Prediction.** To investigate whether orofacial dynamics encode brain-wide neural signals, a feed-forward neural network was trained using 24 input features (x/y coordinates) derived from face-tracking. A preprocessing module enhanced temporal and spatial continuity of these signals before passing them into multi-layer networks (5 or 10 hidden layers). The network output was a predicted trace of calcium activity for each of the 11 brain regions. Model training and testing were performed on GCaMP6s animal data, demonstrating that high-dimensional facial movements carry rich neural information. **(C) Representative Predictions from Different Models.** DeepFace-enabled predictions show strong alignment with recorded brain signals across regions, far surpassing predictions made using the Facemap Refined and Basemodel pipelines. Data shown for a GCaMP6s mouse illustrate the model’s fidelity across motor, sensory, and visual cortices. This example underscores DeepFace’s robustness and precision in decoding brain dynamics from orofacial behaviors. **(D) Cross-Line Model Evaluation.** Coefficient of Determination (R²) was used to quantify brain activity prediction accuracy across all models and transgenic lines. DeepFace consistently outperformed competing models (Facemap Refined and Base) across all five animals in each genotype. The bottom-right panel aggregates prediction accuracy across all conditions and demonstrates DeepFace’s superior generalization and neural decoding capacity (GCaMP6s: p < 0.0001 vs Refined and Basemodel, GCaMP6f: p < 0.05 vs. Refined, p < 0.0001 vs. Basemodel, jGCaMP8m: p < 0.001 vs. Refined, p < 0.0001 vs. Basemodel, paired t-test).

We next tested whether these facial features—extracted by three pipelines (DeepFace, Facemap Basemodel, and Facemap Refined)—could predict neural activity across 11 brain regions using feedforward neural networks (**Fig. 2B**). Each transgenic line had a dedicated model trained on 25 experiments and tested on five withheld sessions. Multiple network architectures were evaluated (5 or 10 hidden units, 50–300 epochs), and the best-performing model per method was selected using coefficient of determination (R²) on test data **(Supp. Fig. 5, Supp. Table 1**). Across all regions and genotypes, DeepFace-based models outperformed those trained on Facemap-derived inputs. In a representative GCaMP6s mouse (**Fig. 2C**), DeepFace yielded high-fidelity predictions that closely tracked true neural signals, while Facemap predictions deviated significantly. This performance advantage held across regions including motor (MOp, MOs), somatosensory (SSp_ll, SSp_ul, SSp_tr), retrosplenial (RSPagl, RSPd, RSPv) and visual (VISa, VISam, VISpm) cortices. Population-level results confirmed the robustness and generalizability of DeepFace-based prediction (**Fig. 2D**). DeepFace achieved significantly higher R² scores than both Facemap Refined and Basemodel across all three lines (**GCaMP6s**: p < 0.0001 vs Refined and Basemodel, **GCaMP6f**: p < 0.05 vs. Refined, p < 0.0001 vs. Basemodel, **jGCaMP8m**: p < 0.001 vs. Refined, p < 0.0001 vs. Basemodel).

These results demonstrate that DeepFace effectively captures rich behavioral features that are tightly coupled to cortical dynamics, enabling accurate, large-scale, and noninvasive brain decoding. Building upon the strengths of existing tools such as DeepLabCut and Facemap, DeepFace is designed to address specific challenges in high-throughput applications—such as cross-session generalization and reduced need for manual retraining. By supporting flexible, model-free inference across diverse experimental conditions, DeepFace advances the scalability of behavioral-neural coupling analysis. This approach not only enables robust brain-state decoding from facial features but also contributes to the growing foundation for noninvasive, video-based biomarkers in both preclinical and translational neuroscience.

## Methods

### Animals

All experiments were conducted in accordance with the guidelines of the Cleveland Clinic Institutional Animal Care and Use Committee (IACUC). Mouse genotyping was performed on ear punches by TransnetYX (Cordova, TN) using their automated real-time PCR platform. Mice were maintained on a 14:10 light: dark cycle with ad libitum access to food and water. The room temperature was regulated between 18 °C and 26 °C. Male and female littermates were separated by sex after weaning and housed with same-sex littermates. Reporter mice were generated by crossing either TIT2L-GC6s-ICL-tTA2)-D (stock no. 031562, Jackson Laboratory), Ai148(TIT2L-GC6f-ICL-tTA2)-D (stock no. 030328, Jackson Laboratory), or TIGRE2-jGCaMP8m-IRES-tTA2-WPRE (stock no. 037718, Jackson Laboratory) with B6.Cg-Tg(Camk2a-cre)T29-1Stl/J mice (CaMKIIα-Cre, stock no. 005359, Jackson Laboratory).

### Surgical Procedures

Surgical procedures closely followed the procedure previously used for whole-cortex widefield imaging^15^. Mice were anesthetized with isoflurane (induction 2.5%; maintenance 1-1.5%). Buprenorphine (3.25 mg/kg), Meloxicam (5 mg/kg), and sterile saline (0.05 mL/g) were administered at the start of surgery. Anesthesia depth was confirmed by toe pinch. Hair was removed from the dorsal scalp (Nair, Vetiva Mini Hair Trimmer) and the area was disinfected with 3 alternating applications of betadine and 70% isopropanol. Bupivacaine (5 mg/kg) was then injected under the skin for local anesthetic before the scalp was removed. The skull was then exposed, cleaned, and dried. The remaining outer skin was affixed in position with tissue adhesive (Vetbond, 3M) for clean surgical margins. We created an outer wall using dental cement (C&B Metabond, Parkell) while leaving as much of the skull exposed as possible. A custom circular headbar (eMachineShop) was secured in place using dental cement (C&B Metabond, Parkell). A layer of optical glue (Norland Optical Adhesive NOA 81, Norland Products) was then applied to the exposed skull and cured with a UV flashlight (LIGHTFE, UV301Plus-365nm). The mice were allowed to fully recover in a warmed chamber and then returned to their home cages. Post-operational care consisted of three daily injections of Meloxicam (5 mg/kg) following the surgery.

### Widefield Imaging of Brain Activity

For wide-field imaging, cortical excitation was performed sequentially using blue (M470L5, Thorlabs) and violet (M405LP1, Thorlabs) light emitting diodes (LEDs). Blue illumination (470 nm) targeted genetically-encoded calcium indicators (GECI) (GCaMP6s, GCaMP6f, and jGCaMP8m), while violet (405 nm) illumination immediately followed to provide a reference measurement used to correct for hemodynamic artifacts. The two excitation wavelengths were combined through a dichroic beamsplitter (87-063, Edmund Optics) and sent onto the cortical surface via an objective lens (MVL50M23, Navitar). Emitted light was collected through an emission dichroic beamsplitter (T495LPXR, Chroma), and filtered using a band-pass filter (86-963, Edmund Optics). The filtered fluorescence signal was then focused on a CMOS camera (CS505MU1, Kiralux) through a second objective lens (MVL16M23, Navitar). Data was acquired at a resolution of 1200 x 1200 pixels with 4×4 pixels binning, resulting in an effective 300×300 pixels covering a cortical area of 9.85 x 9.85 mm^2^. This configuration enabled an acquisition rate of 40 frames per second (fps), sufficient for resolving the kinetics of the GCaMP6s and GCaMP6f calcium indicators. Experiments were carried out at 40 Hz frame rate for blue and violet light combined (effective frame rate of 20 Hz). Average laser powers were maintained below 10mW for the blue LED and below 50 mW for violet LED. Beam diameters were measured by the knife-edge test method, (1/e^2^ criterion) were 9.2 for blue LED and 11.7mm for violet LED. For functional imaging with jGCaMP8m mice, we used another objective (MVL100M23, Navitar) and utilized 1200×1200 pixels with binning 2×2 pixels (effectively 600×600 pixels) which cover 9.85 x 9.85 mm^2^ surface area on the mouse brain. With this configuration, we can achieve 100 frames per second (fps) acquisition rate, which is fast enough to record jGCaMP8m calcium indicator. We used < 10 mW average power for blue LED. By performing the knife-edge test, we measured 1/e^2^ lateral spot sizes of the blue beam as 9.2 mm.

### Imaging of Pupil and Orafacial Features

We recorded the pupil and face of the animal in all mouse lines. To illuminate the left side of the face we used an infrared light source (LIU780A, Thorlabs). The pupil images were collected with a focusing telecentric lens (58-430, Edmund Optics) and a monochromatic camera (1500M-GE-TE, Thorlabs) with a frame rate of 40Hz. We utilized 1000×1000 pixels with binning 4×4 pixels (effectively 250×250 pixels) which cover 9.85 x 9.85 mm^2^ surface area on the mouse pupil. The face images were collected with a separate camera (CS505MU1, Thorlabs) and infinity corrected objective lens (MVL25M23, Navitar) with a frame rate of 120Hz. We utilized 1200×1200 pixels with binning 3×3pixels (effectively 400×400 pixels) which cover 9.85 x 9.85 mm^2^ surface area on the mouse pupil.

### Spontaneous Behavior

Spontaneous locomotor activity was monitored during imaging experiments using a spherical treadmill apparatus designed to quantify voluntary movement in a head-fixed mouse. The treadmill consisted of a 20-cm polystyrene foam ball positioned within a hemispherical 3D printed polylactic acid (PLA) bowl, modified from previous studies^16,17^. The bowl contained eight airflow ports at its base to supply pressurized air, facilitating near-frictionless rotation of the foam ball. Movements were recorded simultaneously in three axes: anterior-posterior (forward-backward), medial-lateral (left-right), and yaw (rotational) using two USB optical motion detectors (Gaming Mouse G302, Logicool) positioned orthogonal at the equator of the treadmill. Mice were head-fixed via a headplate secured to a mounting bar and positioned ∼1 cm above the top of the ball. Mice underwent three days of training to acclimate to head-fixation and the testing environment prior to imaging sessions. On the first day, the animal was head-fixed and was allowed to freely move on the treadmill without airflow. On the second and third days, the pressurized air gradually increased to quarter and half of the final pressure level (70 psi), respectively.

### Allen Brain Atlas Registration

We used the alignment script provided by the locaNMF toolbox^18^ to rigidly align the skull surface’s anatomical landmarks to a standard template atlas. For each animal, we selected seven key points: the left, center, and right intersections between the anterior cortex and olfactory bulb, the location of the bregma, and the base of the retrosplenial cortex on the midline skull suture.

### Hemodynamic Subtraction

For hemodynamic correction, we first computed the blue and violet ΔF/F for each brain region by subtracting and dividing by the median average brain activity during the whole session (5 min). We used linear regression to regress the violet time series on the blue time series, i.e. find coefficients m and b of the regression b = mv + b where b is the blue and v is the violet ΔF/F signal. The subtracted signal was given by b – (mb + b). This gives a single time series per region, which was used for subsequent analyses.

### mediaGUI Methodology

We developed a graphical user interface (GUI) tool to streamline video data preprocessing for training pipelines (**Supp. Fig. 2**). The software allows users to efficiently compress, convert, and concatenate experimental video data through an intuitive interface, minimizing manual effort and significantly improving rendering speed. The current version (v1.0.3) supports uniform frame extraction from a user-specified list of videos, concatenating the frames into either MP4 or AVI outputs.

The tool is a cross-platform and has been successfully tested on Windows 10/11, MacOS 14/15, and Ubuntu 22.04. It is implemented in Python and uses PyQt6 (v6.6.0) for GUI development. Video processing operations are powered by OpenCV-Python (v4.8.1) and numpy (v1.26.0). Where available, the CUDA Toolkit is utilized to offload frame reading and resizing to the GPU, accelerating processing times. On Windows systems, the openh264 library is integrated to enhance MP4 rendering speed, resulting in faster export compared to AVI format.

The application was designed with two primary goals: (1) to provide an intuitive and minimalistic interface for video processing, and (2) to optimize data processing speed. The user interface consists of a drag-and-drop list for video selection, customizable export settings, and a one-click export button for generating the concatenated output. Features and settings are deliberately kept minimal to ensure a streamlined workflow. To optimize performance, if a CUDA-enabled GPU is detected, frame reading and resizing are performed on the GPU. If only a CPU is available, the software applies a combination of batch processing and sequential frame reading strategies to maximize efficiency during even frame extraction. The software is open-source and available at https://github.com/khicken/mediaGUI/, under the MIT License. Installation is recommended through a direct executable download but is also available via pip package installation. The repository provides detailed installation and usage instructions.

### sleapGUI Face Methodology

We developed a graphical user interface (GUI) tool to automate SLEAP processing workflows for animal tracking videos (**Supp. Fig. 2**). The application enables researchers to apply DeepFace models and process tracking data through an intuitive interface, eliminating the need for direct command-line interaction. Unlike the default SLEAP GUI, which requires manual input at multiple steps for each video, our developed tool automates the analysis process, making it significantly easier and faster for users to process large datasets with minimal intervention. The application applies the same analysis parameters used during both local computer and high-performance computing (HPC) processing, ensuring consistency and reproducibility across different platforms. Importantly, the tool is designed to mimic an HPC-style batch analysis workflow on a local computer. By automatically analyzing all selected videos in sequence without requiring user supervision, the GUI enables researchers to efficiently handle large datasets even without access to HPC resources. This design ensures that users working on standard desktop or laptop machines can perform large-scale analyses with the same automation and reliability typically achieved in HPC environments.

The software streamlines three primary tasks: (1) analyzing multiple videos with a selected DeepFace model, (2) exporting tracking data to CSV format for further analysis, and (3) generating labeled visualization videos from analysis results.

The tool is a cross-platform and has been tested on Windows, MacOS, and Linux. It operates within environments configured with Python (v3.7), SLEAP (v1.4.1), and Anaconda (v25.3.0). To ensure efficient video analysis, the local machine must have an NVIDIA GPU with CUDA support, as required by the SLEAP environment for model training and inference. The application is built in Python, leveraging the SLEAP environment’s pre-installed QtPy (v2.4.2) for GUI development. Core functionality relies on DeepFace’s command-line interface, with the GUI acting as an abstraction layer. Threading is employed to maintain a responsive interface during computationally intensive tasks, and platform-specific optimizations ensure reliable output collection across both Windows and UNIX-based systems.

The application was developed with two primary objectives: (1) to provide a streamlined interface that automates common DeepFace workflows, and (2) to ensure robust and reliable processing of long-running video analysis tasks. The user interface features a model selection dropdown (including support for pretrained models), input and output path configuration fields, and clearly labeled action buttons. The application employs as worker-thread system for executing SLEAP commands. The worker class manages subprocess execution with timeout detection, real-time progress reporting, and error handling. Additionally, the GUI limits continuous processing to a maximum of 24 hours per session to ensure system stability during extended analyses while also preventing GPU overclocking or overheating.

The software is open-source and available at https://github.com/khicken/sleapGUI/, under the MIT License. Installation is recommended via pip after activating the SLEAP environment, which installs a console script entry point (sleapgui or sleapgui face) that allows users to launch the application directly from the SLEAP environment command line. Detailed installation and usage instructions are provided in the repository.

### DeepFace and DeepLabCut Methodology

DeepFace utilized SLEAP version 1.3.3, while DLC relied on version 2.3.10 as the backbone for pose estimation. The training video was generated using mediaGUI, a Python-based GUI, by evenly extracting 330 images from each video (90 videos total) coming from GCaMP6s, GCaMP6f, and jGCaMP8m animals that went through either 20 Hz (GCaMP6s and GCaMP6f) or 100 Hz (jGCaMP8m) brain imaging. These videos spanned 15 animals across three different transgenic lines, with each animal undergoing six experiments. Each experimental video contained approximately 36,000 frames. The extracted images were then combined into a single concatenated video (**Fig. 1a and Supp. Fig. 1a**). Once the concatenated video was created, it was inserted into the DLC code package, where 1800 images were evenly extracted for labeling (**Supp. Fig. 1a**). The model consisted of 12 labeled points: 4 on the eyelid, 2 on the nose, 2 on the mouth, and 4 on the whiskers (**Fig. 1a**). To ensure precise labeling, the label size was set to 8 instead of the default 12. The DLC labels were then transferred into the SLEAP code package for DeepFace training. Before training, we defined relationships between different points. The connections were established as follows: top and bottom eyelid, right and left eyelid, top and bottom mouth points, top left and top right whiskers, and bottom left and bottom right whiskers (**Fig. 1a**). The DeepFace model was trained using a Single Animal Training Model with UNet as the backbone. To ensure the model training window covered the entire video, both maximum stride and filters were set to 64, while all other parameters remained at their default settings. Optical Flow, Centroid, and Greedy were used as optimal parameters for data analysis. Before analysis, the 12 labeled points used for training the model were specified.

The DLC model was also trained using the following parameters: a model shuffle of 1, the ResNet-50 architecture, imgaug parameter used for data augmentation to create the ResNet-50 model, a maximum of 1,000 training iterations, checkpoints saved every 500 iterations, progress displayed every 50 iterations, a total of 5 recorded snapshots, and a shuffle value of 1. Once training was completed, the data was inserted into the model for analysis.

The results from both DLC and DeepFace on each original experimental video were saved as CSV files for post-processing analysis (**Supp. Fig. 1a** and **Supp. Fig. 2**). We also developed a custom-built GUI (sleapGUI Face) inside the new SLEAP 1.4.1 environment for users to efficiently analyze the face data, save it as a csv file, and create labeled videos by using the same parameters as we used to analyze the face data in SLEAP version 1.3.3.

### Facemap Methodology

A flowchart was created to illustrate the workflow and demonstrate how Facemap was applied in the analysis (**Figure 1b and Supp. Fig. 1b**). The base model in the Facemap code package was the pretrained model with their own dataset therefore it requires to be retrained with any new data. To improve accuracy of the basemodel, we refined the base model of Facemap for each video using the default graphical user interface (GUI) parameters: 15 images, 50% random frames, and a 95% difficulty threshold. A total of 11 key points were selected for labeling in Facemap, including four eye locations, two nose points, two mouth points, and three whisker points. Four additional points (nose (R), nose (tip), nose (bridge), and paw) were excluded from the refinement process to maintain consistency with the DeepFace and DLC models. Once the model was refined and retrained, the updated version was reapplied to the video, and the output h5 file was converted into CSV format for post-processing analysis using a MATLAB script (**Supp. Fig. 1b**). The refinements were saved for each input model, allowing for progressive improvements in accuracy over time. This process was repeated for each video to enhance the model’s precision and reliability.

### High Performance Computing Methodology

High-Performance Computing (HPC) was utilized for DeepFace analysis to process all videos in a high-throughput manner. This was achieved by running a single SLURM job, which leveraged parallel computing to accelerate inference speed (**Supp. Fig. 2**). A batch SLURM script was created and submitted to the HPC to analyze all videos using SLEAP’s command-lines. The analysis was conducted on an A100 chip with the following parameters: 2 nodes, 3 GPUs per node, and 128 GB per GPU.

### Brain Activity Prediction using custom Deep Learning Model

We developed a MATLAB based GUI application to build and evaluate a neural network model that predicts brain signal at time (t) from mouse facial movement features. The primary analytical steps involve data acquisition, preprocessing and feature extraction, temporal alignment of modalities, neural network training, and comprehensive evaluation using multiple performance metrics.

Facial movements were recorded at 120 Hz with our custom-made system and analyzed using either DeepFace or FaceMap software, resulting in CSV files containing the raw two dimensional *(x,y)* coordinates of facial landmarks, specifically the eyes, nose, mouth, and whiskers. Concurrently, neural activity was recorded using hemodynamic-subtracted widefield calcium imaging from 11 cortical regions in both hemispheres, sampled at 20Hz in GCaMP6s and GCaMP6f transgenic mouse lines and at 100 Hz in jGCaMP8m mice. Since we recorded the facial movements on the left side, we included the activity of 11 brain regions on the right hemisphere. Each facial recording (CSV) was systemically paired with its corresponding brain activity recording (.mat file) from 11 different regions to create matched face-brain data pairs for subsequent model training and evaluation.

The raw facial coordinate data underwent multiple stages of preprocessing to extract meaningful spatial and dynamic features from each recorded frame. Spatial features were computed as follows: eye features were derived by calculating the centroid from the four points of the eyes; nose and mouth features were obtained by calculating the midpoint coordinates between their respective top and bottom landmarks. For example, the nose midpoint coordinates (x_nose_, y_nose_) were computed as:

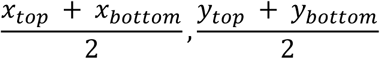

Whisker features were similarly computed as the centroid of four points for DeepFace or three points for FaceMap recordings, calculated as the mean position across whisker coordinates. Following spatial feature extraction, velocity features were derived by computing the frame-to-frame spatial differences multiplied by the sampling rate (120 Hz), thereby capturing the dynamic aspects of facial movement over time. The resulting 12 spatial and 12 velocity features (11 velocity features in FaceMap) were combined into a 24-dimensional raw feature vector per frame. Missing or invalid data points were subsequently removed and replaced using linear interpolation with end-extrapolation. For temporal dynamics, a sliding-window aggregation of five consecutive frames was performed. This reshaped each sequence of 5 consecutive 24-dimensional vectors into one flattened 120-dimentional vector.

Neural activity signals were extracted from hemodynamic-subtracted fluorescence signals for GCaMP6s and GCaMP6f and raw fluorescence signals for jGCaMP8m. These signals were band-passed filtered using a second-order Butterworth filter with cutoff frequencies at 0.1 and 9.99Hz for 20Hz imaging and 0.1 to 49.99 for 100Hz imaging to isolate physiologically relevant signal frequencies and to remove drifts. Filtering was performed in both forward and backward directions using MATLAB’s “*filtfilt*” function to ensure zero-phase distortion. Sequentially, signals were normalized by computing the ΔF/F. To reduce high-frequency fluctuations, ΔF/F signals were smoothed using a 1-s moving-average filter.

To enhance predictive power, we also added principal component (PCA) analysis across the 11 brain regions, and the first three principal components were retained. Derived brain features were linearly interpolated to exactly match the duration of the processed brain activity data. The first 5 seconds of data from both modalities were removed to eliminate initial recording transients and filter artifacts. Subsequentially, facial features (originally at 120Hz) were interpolated to match the brain activity timestamps (20 Hz or 100 Hz, depending on mouse strain). Derived PCA-based brain features were also interpolated to the brain timestamps. For training, an auto-regressive lag feature was included by incorporating the previous timepoint’s brain ΔF/F signal as an additional predictive feature. Thus, at each timepoint, t, the complete feature vector was constructed as:

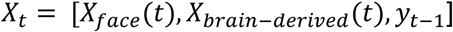

Training pairs {X*^(i)^*, Y*^(i)^*}^12^ were vertically concatenated into comprehensive matrices:

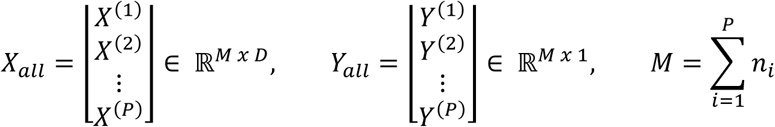

A feed-forward neural network model with one hidden layer (tested at 5 and 10 neurons) was used. Hidden layer neurons used hyperbolic tangent activation functions, and the output layer was a linear neuron predicting *y_t_*. Model parameters were optimized via MATLAB’s Levenberg-Marquardt backpropagation method “*fitnet*,” approximating Gauss-Newton updates. Training minimized mean-squared error (MSE):

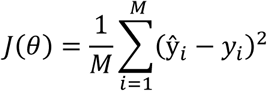

Training concluded upon reaching either the maximum epoch count (50, 100, 250, or 300 epochs) or when MSE fell below 10^-4^. Model performance metrics calculated included root mean square error (RMSE), coefficient of determination (R^2^), and mean absolute error (MAE).

During testing, predictions *y*^ were made on independent datasets. Performance was quantified similar to training, calculating RMSE, MAE, and R^2^:

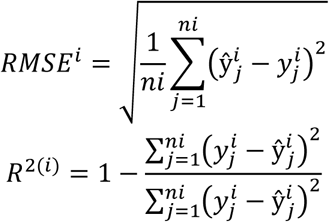

Also, the actual (*y*^*i*^) and predicted (ŷ^*i*^) brain signal is plotted, where the overall performance is summarized by computing the mean and standard deviation of the *RMSE*^*i*^ and *R*^2(*i*)^ of all test pairs.

## Acknowledgements

This work was supported by US National Institute of Health (NIH) grants R00 EB027706 (MY), Cleveland Clinic and IBM Discovery Accelerator Grant (MY), and Case Western University SOURCE Fellowship (KO). Figures 1a–d, 2a–c, Supplementary Figures 1–2 and 4–5, as well as the GCaMP6s, GCaMP6f, and jGCaMP8m illustrations featured in the Supplementary Videos 1–6, were created using BioRender.com.

## SUPPLEMENTARY FIGURES

**Supp. Fig. 1:**
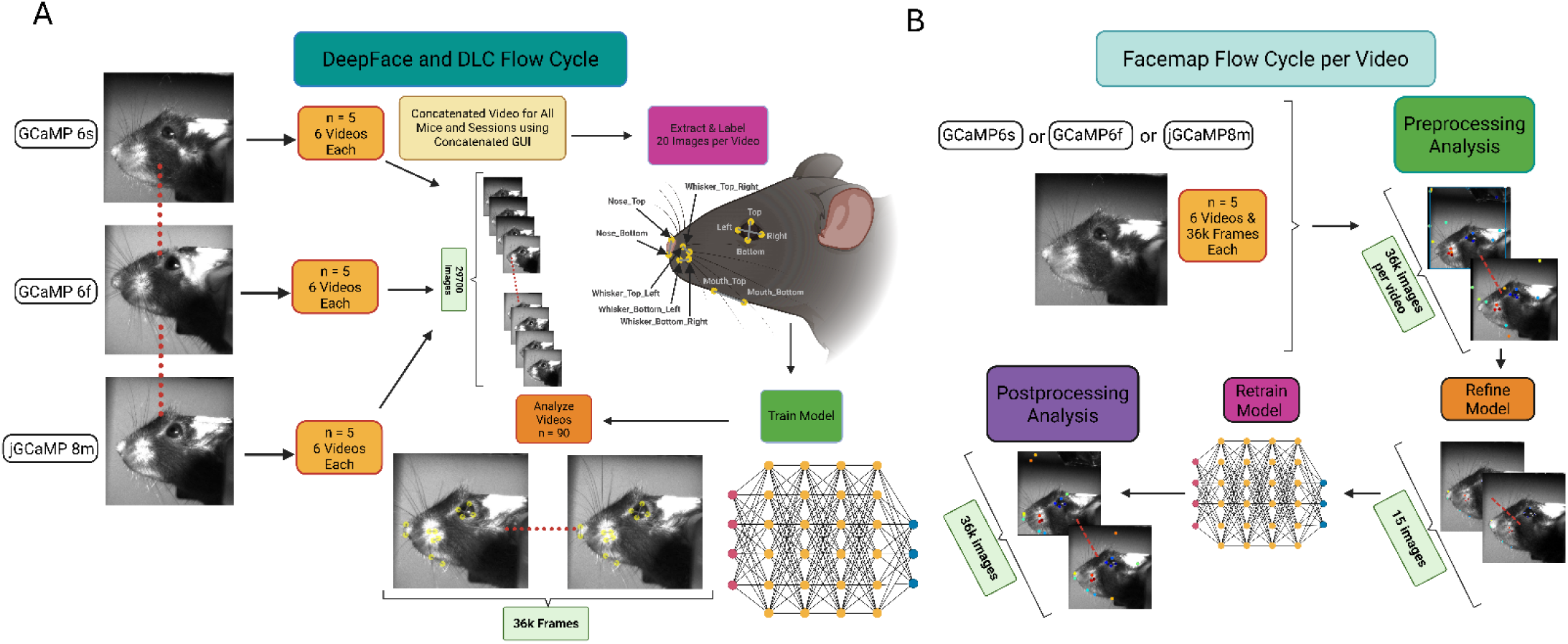
Workflow Comparison of DeepFace/DLC and Facemap Pipelines for Orofacial Feature Tracking in Mice. **(A)** Detailed Clockwise Flow Cycle for DeepFace and DeepLabCut. In the first part, 330 frames were extracted evenly for every single experimental video (n = 90) and combined to create a concatenated video for all experiments using the custom Concatenation GUI. Next, the concatenated video was uploaded to SLEAP and DeepLabCut pose estimation code packages, and 1800 images were extracted for labeling using a manual method. After extracting the frames, the images were labeled using 12 keypoints (4 for eye, 2 for nose, 2 for mouth, and 4 for whisker). Once the images were labeled, the model was trained on UNet Backbone (SLEAP) or ResNet50 (DeepLabCut), and the raw videos were analyzed for each experiment using the custom DeepVision Analysis GUI. **(B)** Detailed Clockwise Flow Cycle for Facemap. An experimental video (n = 90) (Top Left) is uploaded to the Facemap code package. Then (Top Right), the data is analyzed using the base model. Afterwards (Bottom Right), the model is refined by labeling 15 images using 11 keypoints (4 for eye, 2 for nose, 2 for mouth, and 3 for whisker), and then retrained. Finally (Bottom Left), the video is reanalyzed using the refined model.

**Supp. Fig. 2.**
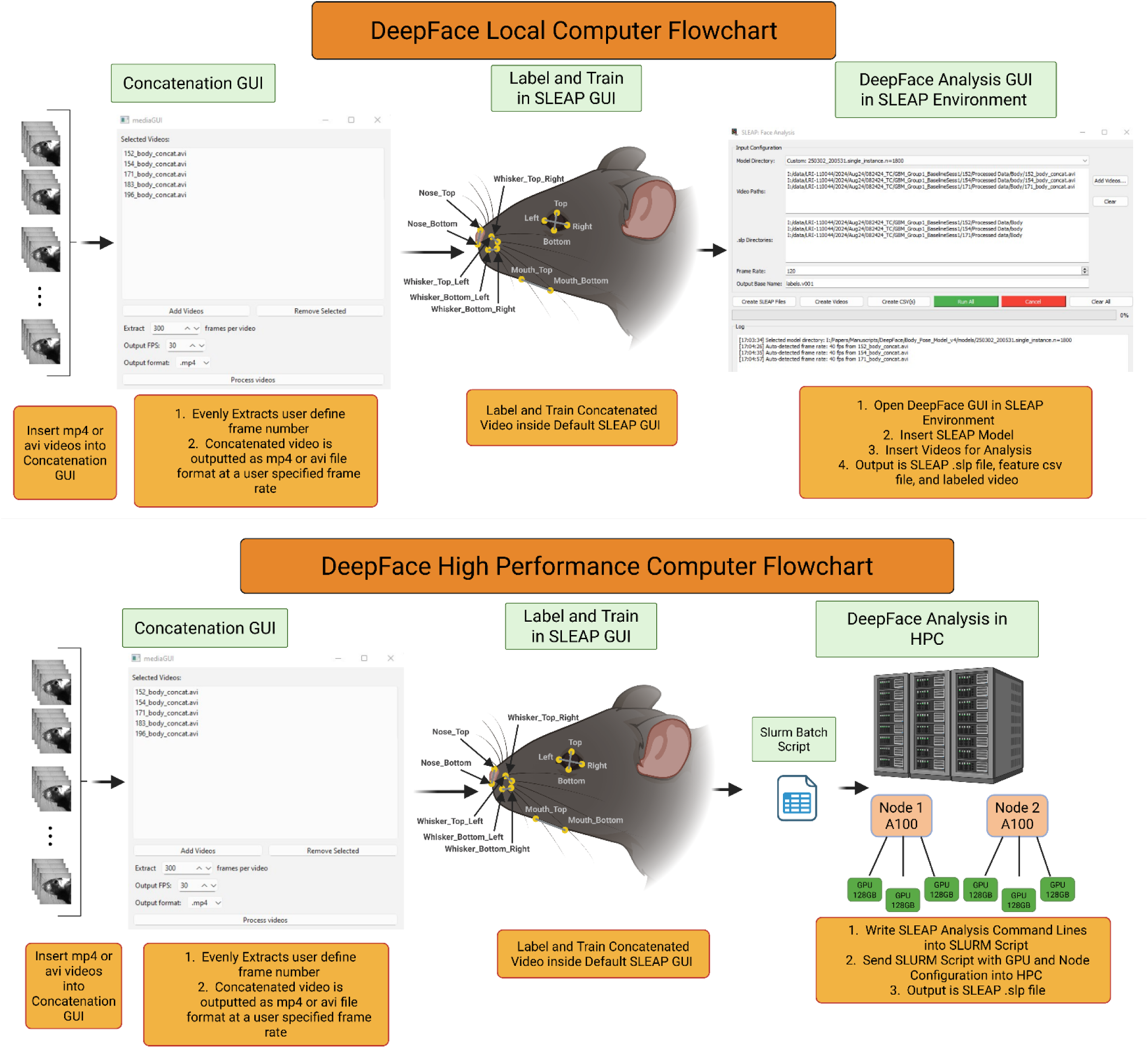
DeepFace Workflow for Local and High-Performance Computing Environments Caption. **(A)** DeepFace Detailed Analysis Flow Chart for Local Computer. All of the experimental face videos (either mp4 or avi video format) are inserted into the Concatenation GUI for creating a concatenated video (mp4 or avi format). The GUI will ask the user to specify the number of frames it will be extracted evenly per video, as well as the concatenated video frame rate and output location. After the video is concatenated, it is uploaded into the SLEAP default GUI for labeling and training the model. After training the model, the custom DeepFace Analysis GUI is opened inside the SLEAP Anaconda environment for analyzing the videos. Once the GUI opens, it requires the user to first input the SLEAP model location, then insert the desired videos into the video path location. After the videos are placed into the video path location, the GUI automatically generates the SLEAP analysis file (.slp), feature coordinates csv file, and labeled video inside the same file path as the inserted raw video. Finally, clicking the Run All green button will analyze all of the inserted video data. **(B)** DeepFace Detailed Analysis Flow Chart for High Performance Computer. The DeepFace analysis process is the exact same as in the local computer until after training the SLEAP model. Once training the model, a slurm batch script is created to analyze the video data. Inside the script, the number of nodes and GPUs are specified, as well as the SLEAP commands to analyze all of the videos. Then, the slurm batch script is sent to the HPC for video analysis and the output is the SLEAP analysis file.

**Supp. Fig. 3.**
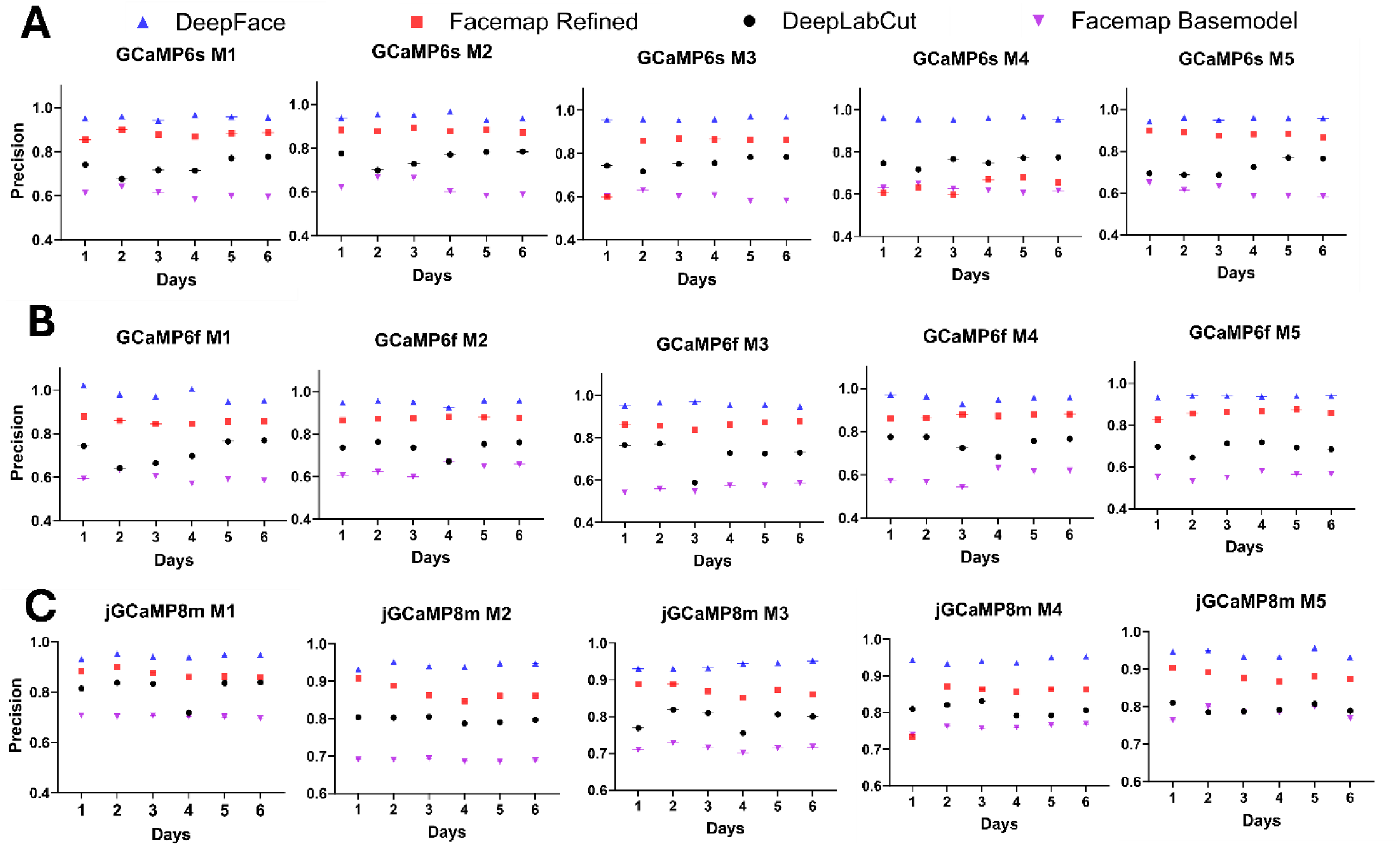
DeepFace Outperforms Existing Methods in Orofacial Feature Tracking Across GCaMP Mouse Lines Over Multiple Days. **(A)** Detailed precision values for GCaMP6s animals. In each mouse (n = 5) across six experimental days, the precision and standard error was plotted for each day on y axis with respect to each model—DeepFace, Facemap Refined, DeepLabCut (DLC), and Facemap Base Model. The same detailed precision values were done for **(B)** GCaMP6f and **(C)** jGCaMP8m animals.

**Supp. Fig. 4.**
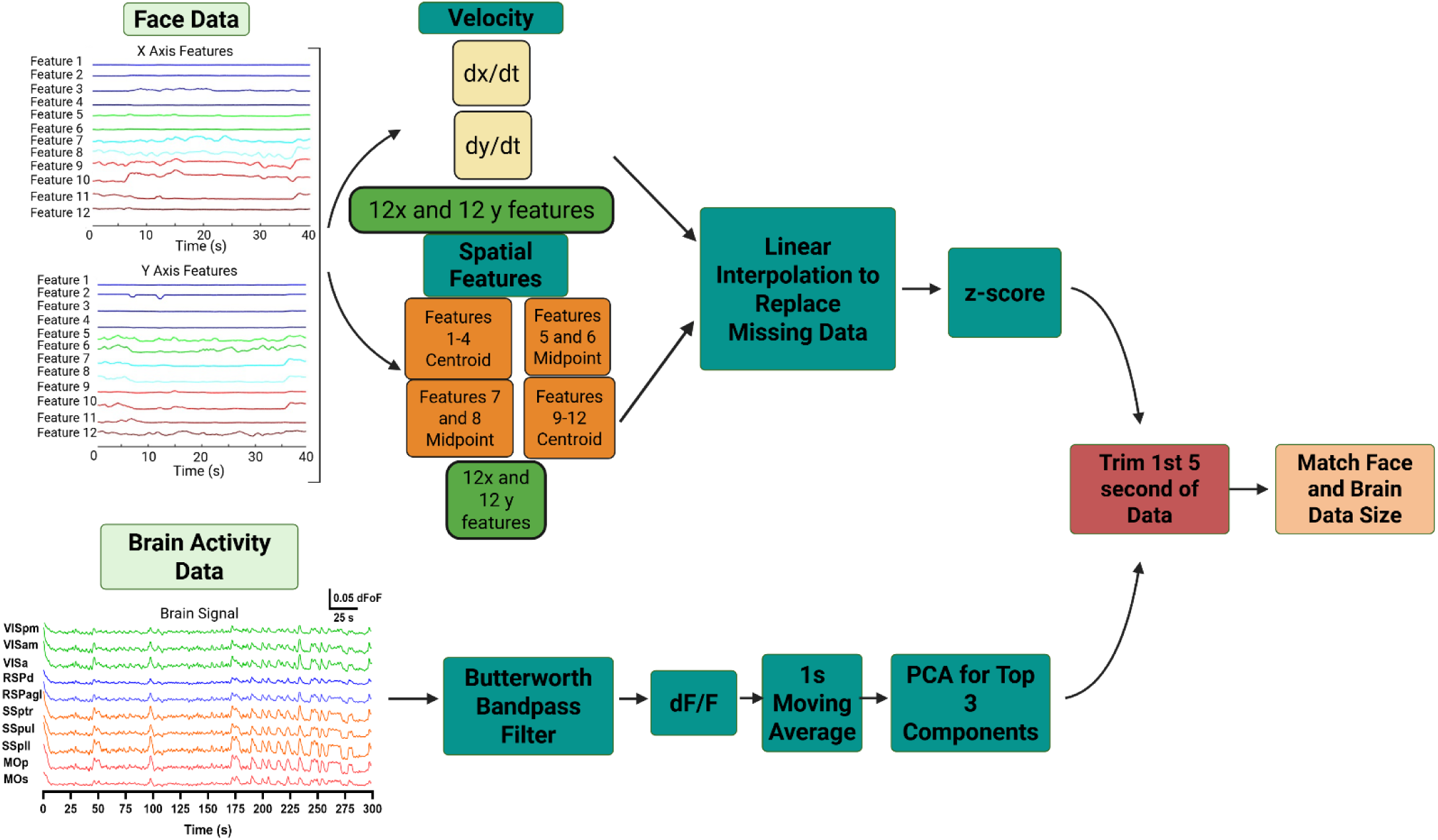
Preprocessing Pipeline for Facial and Brain Activity Data Integration Caption. Detailed Facial Feature and Brain Activity Processing for Brain Activity Model. The facial features from each animal and each of the trained days (1, 2, 3, 5, and 6) underwent the processing technique. The x and y features were separated into two components, where in the 1^st^ component, the derivatives of the features were taken. In the second component, spatial features were clustered together by finding the centroid for the eyelid features (1 through 4) and whisker features (9-12), and the midpoint for the nose (5 and 6) and mouth (7 and 8) features. Afterwards, both components underwent linear interpolation to replace the missing data and were z-scored. The linked brain activity, meanwhile, went through Bandpass filter using Butterworth. Then dF/F was applied to the brain, then 1 second moving average, then PCA was done for top 3 components. Afterwards, the brain data merged with the face features, where the first 5 seconds of both data were trimmed to avoid initial recording affects along with filter/transient artifacts. Finally, the 120 Hz face data was reduced dimensionally to match the brain activity data (20Hz for 6s and 6f and 100 Hz for 8m) to feed into the model.

**Supp. Fig. 5:**
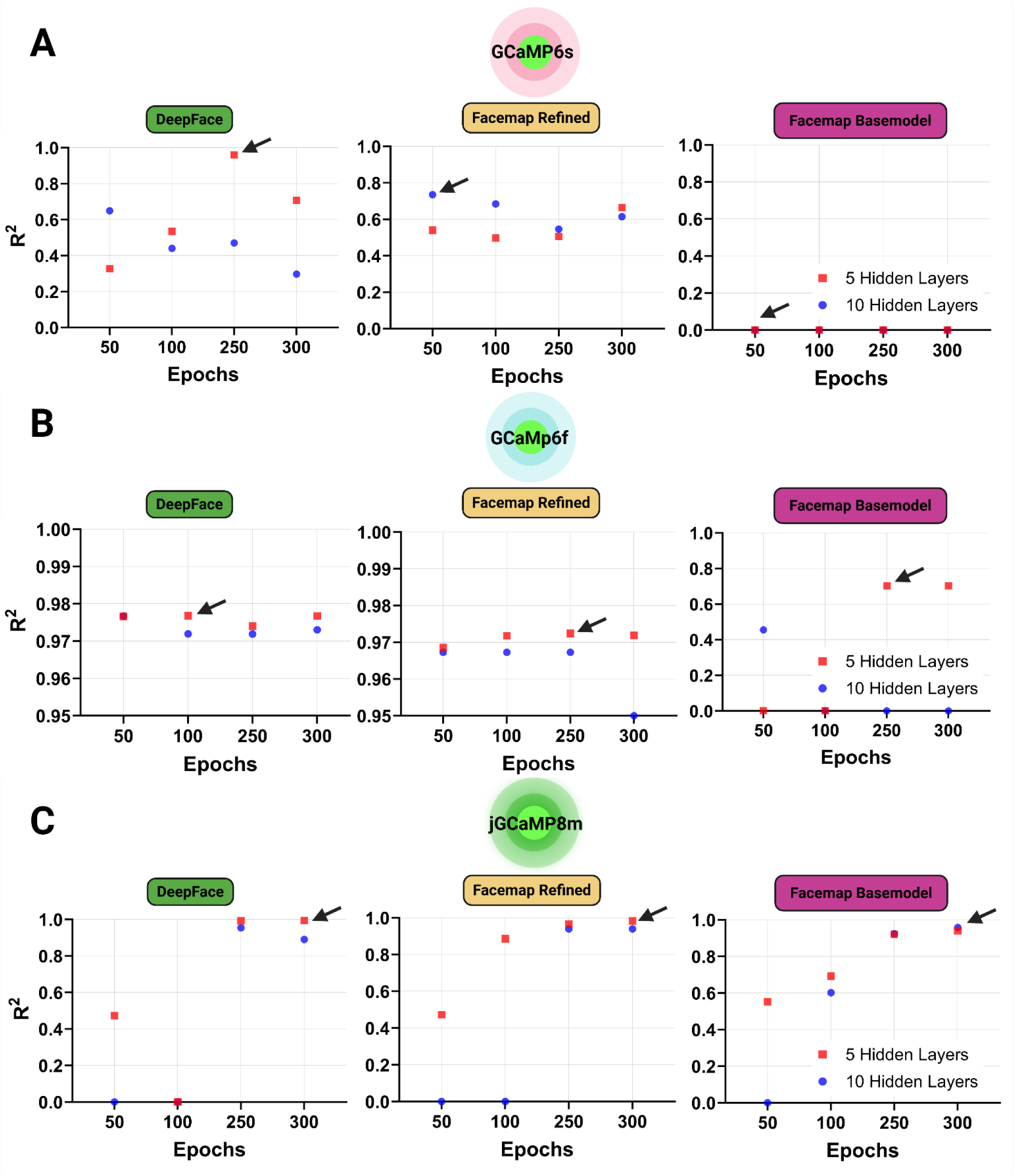
Comparison of brain activity prediction accuracy (R²) using orofacial features extracted by DeepFace, Facemap refined, and Facemap base models across GCaMP6s (A), GCaMP6f (B), and jGCaMP8m (C) mouse lines. For each model, prediction performance is shown across different training epochs (50–300) and network configurations (5 hidden layers: red squares; 10 hidden layers: blue circles). DeepFace consistently achieves high R² values with minimal variability across training conditions, Arrows indicate the best-performing configuration for each method and condition.

**Supplementary Table 1.** Comparison of predictive performance of facial motion based deep learning models (DeepFace, Facemap base model, and Facemap refined model) across three genetically encoded calcium indicator lines (GCaMP6s, GCaMP6f, and jGCaMP8m). Facial motion data extracted using either DeepFace or Facemap tools were used as inputs for a feedforward neural network deep learning model trained to predict neuronal calcium activity. The model was trained separately for each calcium indicator using data from five animals per indicator line, with six experiments conducted per animal. For each animal, five experiments were utilized for model training and one recording was withheld as a testing dataset. Models were systemically trained with various hyperparameter combinations, specifically varying the number of hidden neurons (5 or 10) and training epochs (100, 250, or 300 epochs). The optimized parameters for each condition were determined based on the combination that yielded the highest coefficient of determination (R^2^) and lowest Root Mean Square Error (RMSE) on the held-out testing dataset. The performance metrics (RMSE and R^2^) reported here represent the optimized results, which were subsequently used for all further analyses and graphical representation presented in the main manuscript.

|  | **GCaMP6s** |  |  |  | **GCaMP6f** |  |  |  | **jGCaMP8m** |  |  |  |
| --- | --- | --- | --- | --- | --- | --- | --- | --- | --- | --- | --- | --- |
| | Hid Neuron | Epoch | RMSE | $R^2$ | Hid Neuron | Epoch | RMSE | $R^2$ | Hid Neuron | Epoch | RMSE | $R^2$ |
| DeepFace | 5 | 250 | 90.516 | 0.959 | 5 | 100 | 78.06 | 0.977 | 5 | 300 | 19.245 | 0.999 |
| Facemap Base Model | 5 | 50 | 241.333 | 0 | 5 | 250 | 104.74 | 0.708 | 10 | 300 | 49.364 | 0.958 |
| Facemap Refined Model | 10 | 50 | 101.526 | 0.508 | 5 | 250 | 220.13 | 0.972 | 5 | 300 | 25.349 | 0.983 |

## VIDEO CAPTIONS

**Supplementary Video 1.** Comparison of orofacial tracking performance across DeepFace, DeepLabCut (DLC), Facemap base model, and Facemap refined model in a head-fixed GCaMP6s mouse (Mouse 3, Day 1 experiment). The same video is processed by each method, with facial landmarks—including the eyelid, nose, mouth, and whiskers—overlaid on the raw footage. DeepFace demonstrates superior precision and stability during spontaneous facial movements and variable illumination conditions. This video supports the comparative analysis presented in Figure 1C.

**Supplementary Video 2.** Comparison of orofacial tracking performance across DeepFace, DeepLabCut (DLC), Facemap base model, and Facemap refined model in a head-fixed GCaMP6f mouse (Mouse 1, Day 4 experiment). The same video is processed by each method, with facial landmarks—including the eyelid, nose, mouth, and whiskers—overlaid on the raw footage. DeepFace demonstrates superior precision and stability during spontaneous facial movements and variable illumination conditions. This video supports the comparative analysis presented in Figure 1C.

**Supplementary Video 3.** Comparison of orofacial tracking performance across DeepFace, DeepLabCut (DLC), Facemap base model, and Facemap refined model in a head-fixed jGCaMP8m mouse (Mouse 4, Day 1 experiment). The same video is processed by each method, with facial landmarks—including the eyelid, nose, mouth, and whiskers—overlaid on the raw footage. DeepFace demonstrates superior precision and stability during spontaneous facial movements and variable illumination conditions. This video supports the comparative analysis presented in Figure 1C.

**Supplementary Video 4.** Representative example of simultaneous orofacial tracking and brain activity visualization in a head-fixed GCaMP6s mouse (Mouse 3, Day 1 experiment). The top left panel shows cortical activity extracted from widefield calcium imaging, while the top right panel displays facial tracking results from DeepFace, highlighting real-time detection of the eyelid, nose, mouth, and whiskers. The bottom panel presents synchronized traces of orofacial feature movements and primary motor cortex activity over a 30-second period. This video illustrates the temporal coupling between facial dynamics and cortical signals, as analyzed in the main text.

**Supplementary Video 5.** Representative example of simultaneous orofacial tracking and brain activity visualization in a head-fixed GCaMP6f mouse (Mouse 1, Day 4 experiment). The top left panel shows cortical activity extracted from widefield calcium imaging, while the top right panel displays facial tracking results from DeepFace, highlighting real-time detection of the eyelid, nose, mouth, and whiskers. The bottom panel presents synchronized traces of orofacial feature movements and primary motor cortex activity over a 30-second period. This video illustrates the temporal coupling between facial dynamics and cortical signals, as analyzed in the main text.

**Supplementary Video 6.** Representative example of simultaneous orofacial tracking and brain activity visualization in a head-fixed jGCaMP8m mouse (Mouse 4, Day 1 experiment). The top left panel shows cortical activity extracted from widefield calcium imaging, while the top right panel displays facial tracking results from DeepFace, highlighting real-time detection of the eyelid, nose, mouth, and whiskers. The bottom panel presents synchronized traces of orofacial feature movements and primary motor cortex activity over a 30-second period. This video illustrates the temporal coupling between facial dynamics and cortical signals, as analyzed in the main text.

## References

1 Musall, S., Kaufman, M. T., Juavinett, A. L., Gluf, S. & Churchland, A. K. Single-trial neural dynamics are dominated by richly varied movements. Nature Neuroscience 22, 1677-+ (2019). 10.1038/s41593-019-0502-4

2 Stringer, C. et al. Spontaneous behaviors drive multidimensional, brainwide activity. Science 364, 255-+ (2019). ARTN eaav7893 10.1126/science.aav7893

3 Clancy, K. B., Koralek, A. C., Costa, R. M., Feldman, D. E. & Carmena, J. M. Volitional modulation of optically recorded calcium signals during neuroprosthetic learning. Nature Neuroscience 17, 807–809 (2014). 10.1038/nn.3712

4 Pitcher, D., Dilks, D. D., Saxe, R. R., Triantafyllou, C. & Kanwisher, N. Differential selectivity for dynamic versus static information in face-selective cortical regions. Neuroimage 56, 2356–2363 (2011). 10.1016/j.neuroimage.2011.03.067

5 Bizley, J. K. & Cohen, Y. E. The what, where and how of auditory-object perception. Nature Reviews Neuroscience 14, 693–707 (2013). 10.1038/nrn3565

6 Gurovich, Y. et al. Identifying facial phenotypes of genetic disorders using deep learning. Nat Med 25, 60-+ (2019). 10.1038/s41591-018-0279-0

7 Bandini, A. et al. Analysis of facial expressions in parkinson’s disease through video-based automatic methods. J Neurosci Meth 281, 7–20 (2017). 10.1016/j.jneumeth.2017.02.006

8 Hsu, A. I. & Yttri, E. A. B-SOiD, an open-source unsupervised algorithm for identification and fast prediction of behaviors. Nature Communications 12 (2021). ARTN 5188 10.1038/s41467-021-25420-x

9. Hernández, E. P., et al. Facial Paralysis Algorithm: A Tool to Infer Facial Paralysis in Awake Mice. Eneuro 12 (2025). Artn 2025 10.1523/Eneuro.0384-24.2025

10 Syeda, A. et al. Facemap: a framework for modeling neural activity based on orofacial tracking. Nature Neuroscience 27, 187–195 (2024). 10.1038/s41593-023-01490-6

11 Mathis, A. et al. DeepLabCut: markerless pose estimation of user-defined body parts with deep learning. Nature Neuroscience 21, 1281-+ (2018). 10.1038/s41593-018-0209-y

12 Pereira, T. D. et al. SLEAP: A deep learning system for multi-animal pose tracking (vol 19, pg 486, 2022). Nature Methods 19, 628–628 (2022). 10.1038/s41592-022-01495-2

13 Chen, T. W. et al. Ultrasensitive fluorescent proteins for imaging neuronal activity. Nature 499, 295–300 (2013). 10.1038/nature12354

14 Zhang, Y. et al. Fast and sensitive GCaMP calcium indicators for imaging neural populations. Nature 615, 884–891 (2023). 10.1038/s41586-023-05828-9

15 Ye, Z. et al. Brain-wide topographic coordination of traveling spiral waves. bioRxiv, 2023.2012.2007.570517 (2025). 10.1101/2023.12.07.570517

16 Buchner, E. Elementary movement detectors in an insect visual system. Biological Cybernetics 24, 85–101 (1976). 10.1007/BF00360648

17 Harvey, C. D., Collman, F., Dombeck, D. A. & Tank, D. W. Intracellular dynamics of hippocampal place cells during virtual navigation. Nature 461, 941–946 (2009). 10.1038/nature08499

18 Saxena, S. et al. Localized semi-nonnegative matrix factorization (LocaNMF) of widefield calcium imaging data. PLoS Comput Biol 16, e1007791 (2020). 10.1371/journal.pcbi.1007791

